# Proposal for practical multi-kingdom classification of eukaryotes based on monophyly and comparable divergence time criteria

**DOI:** 10.1101/240929

**Authors:** Leho Tedersoo

## Abstract

Much of the ecological, taxonomic and biodiversity research relies on understanding of phylogenetic relationships among organisms. There are multiple available classification systems that all suffer from differences in naming, incompleteness, presence of multiple non-monophyletic entities and poor correspondence of divergence times. These issues render taxonomic comparisons across the main groups of eukaryotes and all life in general difficult at best. By using the monophyly criterion, roughly comparable time of divergence and information from multiple phylogenetic reconstructions, I propose an alternative classification system for the domain Eukarya to improve hierarchical taxonomical comparability for animals, plants, fungi and multiple protist groups. Following this rationale, I propose 32 kingdoms of eukaryotes that are treated in 10 subdomains. These kingdoms are further separated into 43, 115, 140 and 353 taxa at the level of subkingdom, phylum, subphylum and class, respectively (http://dx.doi.org/10.15156/BIO/587483). Most of the names have been used previously or these were deduced from those of the type taxa to be able to unambiguously link genera to higher taxonomic levels. In the era of phylogenomics, understanding about the phylogenetic relationships among organisms is rapidly increasing. Classifications systems must keep pace with this race to serve the research community by consistent improvements in precision in terms of taxonomic resolution and maintaining monophyly of the ingredient taxa.

## Introduction

Naming and classification of organisms represents a corner stone for communicating biota and their phylogenetic relationships. Marker gene-based analyses and, more recently, genomics methods have greatly improved our understanding of phylogenetic relationships among biological organisms and enabled phylogenetic classification of many taxonomic groups, for example bacteria and archaea (http://taxonomicoutline.org/), flowering plants (APG 2016), fungi (Spatafora et al. 2016) and multiple groups of protists (Berney et al. 2017). These higher-level classifications and taxonomic treatments of genera, families and orders have been incorporated into general classification systems of NCBI (https://www.ncbi.nlm.nih.gov/taxonomy/), SILVA (Quast et al. 2013) and many others. Several authors have attempted to formalize the classification of life (Adl et al. 2012; Cavalier-Smith 2013; Ruggiero et al. 2015; Drozdov 2017), but all these systems share several common weaknesses (Figure 1). First, these classifications include higher taxa that are intentionally kept paraphyletic due to the paucity of separating morphological characters or very small size of these groups (Figure 1b). Second, many obvious kingdom- and phylum-level groups are only described at the genus or family level, which hampers understanding their actual level of taxonomic distinctness and relative phylogenetic deepness (Figure 1c). Third, taxa at different taxonomic levels may exhibit identical names, which may cause misunderstanding especially when new higher taxa are erected and when scripts are used to assign OTUs to taxonomy (Figure 1d). Fourth, names of higher level taxa do not give a clue to non-systematicists about the ingredient taxa (e.g. supergroup ‘Opisthokonta’, phylum ‘Paramycia’ and class ‘Cristidiscoidea’ containing genera *Nuclearia* and *Fonticula*; Ruggiero et al. 2015)(Figure 1e). The same issue appears to the informal non-Linnaean names such as ‘SAR’ (Burki et al. 2007), ‘LKM11’ (Quast et al. 2013), ‘clade GS01’ (Tedersoo et al. 2017), etc., but in the two latter examples these names were introduced to communicate undescribed taxa. Fifth, some authors generate large amounts of names for most of the nodes in phylogenies, in spite of anticipating that the groups are poorly supported, sometimes paraphyletic, and subject to change in the next analysis with improved taxon sampling and more genetic information (Figure 1f). Sixth, the available classification systems, especially NCBI and SILVA include higher taxa, some of which are separated into >20 taxonomic levels (e.g. Diptera), whereas for many others, only 1–2 levels exist (Figure 1g). For example, a genus may belong directly to a class, which in turn belongs to a kingdom with no intermediate levels. The latter issue is particularly problematic when assigning taxonomy to sequencing-derived ecological data sets. In high-throughput sequencing, tens of thousands of Operational Taxonomic Units (OTUs) commonly require taxonomic assignment, which is usually performed against reference sequence databases based on BLASTn searches, Bayesian classifiers or evolutionary placement algorithms (Bik et al. 2012). Such highly skewed classifications associated with these databases hamper building hierarchical classifications within ecological data sets and may require substantial taxonomic expertise to arrange suitable-level taxonomic groups for comparison (e.g. Bates et al. 2013; Geisen et al. 2015; Bahram et al. 2016). Genera, phyla, orders and classes are the most commonly used taxonomic levels for grouping.

**Figure 1.**
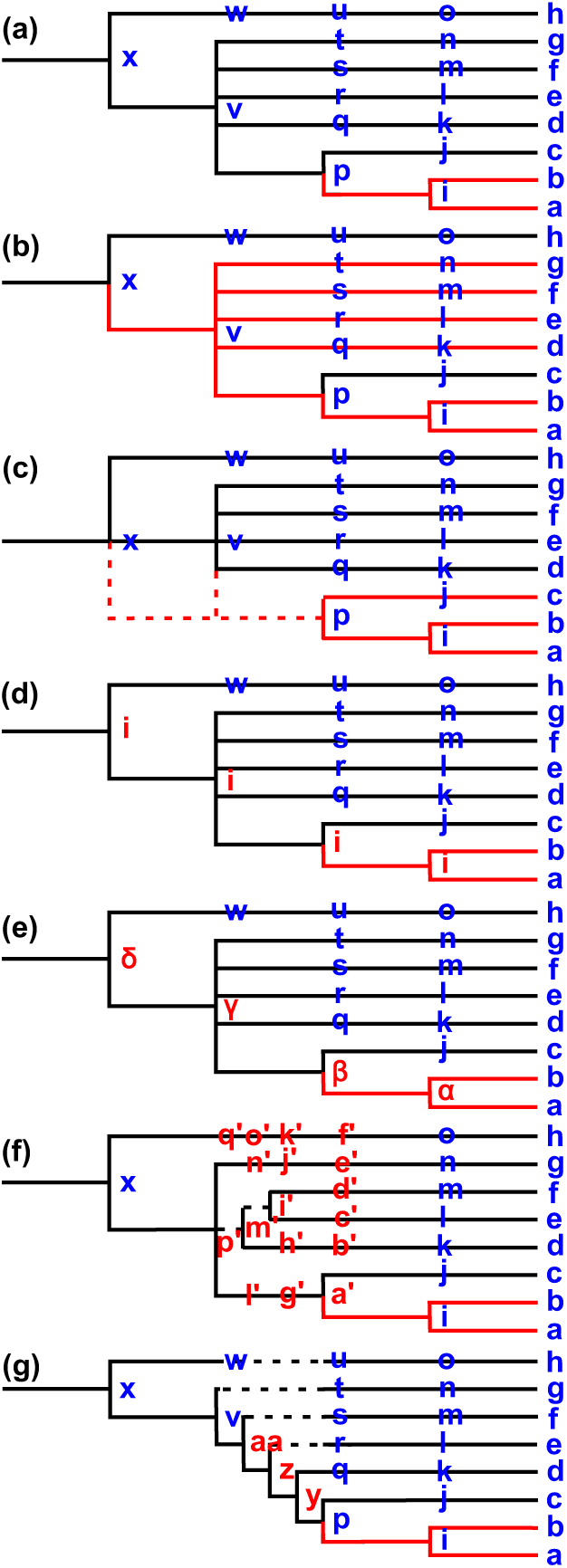
Conceptual scheme indicating major problems in current classification systems. (a) hierarchical classification indicating an example focal group (in red); (b) para- and polyphyletic taxa (clades in red); (c) lack of higher-level resolution and information about taxonomic deepness (clades in dashed line); (d) overlapping names across multiple ranks (names in red); (e) names linguistically unrelated to any ingredient taxon (Greek letters); (f) ephemeral names used for multiple, often poorly supported clades (letters with prim); and (g) differential resolution among taxa in different clades. Letters depict taxon names.

Out of these multiple shortcomings in classification systems, I find the issue of polyphyly and paraphyly the most problematic, because non-monophyletic entities generate taxonomic uncertainty and confusion. Use of e.g. kingdom ‘Protista’ and kingdom ‘Choanozoa’ (sensu Ruggiero et al. 2015) does reflect gross morphology, but provides very limited information about the phylogenetic placement of these groups. Already >50 years ago, Hennig (1966) argued that all taxa should be monophyletic to provide unambiguous understanding of their constituents. Similarly, Avise & John (1999) advocated for the monophyly criterion in classifications based on DNA sequence data and further argued that taxonomic ranks should reflect divergence times to enable comparisons across kingdoms. Decades later we find ourselves still swamped in classification systems comprised of non-monophyletic taxa and finding our way out among multiple synonyms caused by blurring of the boundaries between classical ‘botanical’ and ‘zoological’ systems and attempts to provide short-standing names to nearly each node in the ephemeral phylogenetic reconstructions.

Here I revise the subphylum to subdomain level classification of the Eukarya domain (Woese et al. 1990) focusing on formally described groups and neglecting informal names of ‘environmental’ sequence groups. Based on published molecular phylogenies, NCBI taxonomy backbone, monophyly criterion and comparable level of divergence, I propose a 10-rank alternative classification focusing on subphylum, phylum, subkingdom, kingdom and subdomain levels, with a particular attention to the main ranks above class level. This preprint seeks constructive criticism from the research community to prepare a practical consensus classification of all life that would be efficient for taxon communication among taxonomists and ecologists.

## Methods

Because multiple regularly updated and versioned classifications exist, I first sought to screen the existing systems – UniEuk (www.unieuk.org), NCBI, GBIF (www.gbif.org), SILVA and aforementioned articles - for the best suitable taxonomic backbone. Monophyly of higher-level taxa and use of officially described names were the main criteria for selection. I compared the classifications against >200 phylogenetic studies from class to kingdom levels, giving priority to studies with larger ingroup, greater number of genes and most recent treatments (for minor deep diverging groups). In brief, the following studies were used to extract much of the class to domain level classification information: Yoon et al. (2006), Ruhfel et al. (2014), Magallon et al. (2015), Leliaert et al. (2017) (Archaeplastida); Fiore-Donno et al. (2010), Lahr et al. (2013), Cavalier-Smith et al. (2015b, 2016), Tekle et al. (2016), Kang et al. (2017), Tekle & Wood 2017 (Amoebozoa); Kolisko 2011, Kamikawa et al. (2014), Radek et al. (2014), Cavalier-Smith (2016), Yubuki et al. (2017) (Excavata); Grant et al. (2009), Riisberg et al. (2009), Brown & Sorhannus (2010), Yang et al. (2012), Cavalier-Smith & Scoble (2013), Shiratori et al. (2015, 2017), Yubuki et al. (2015), Aleoshin et al. (2016), Derelle et al. (2016), Dumack (2016), Gao et al. (2016), Krabberød et al. 2017, Reñé et al. (2017) (Harosa); Brown et al. (2009, 2013), Cavalier-Smith & Chao (2010), Zhang (2011), Glücksman et al. (2013), Nosenko et al. 2013, Paps et al. (2013), Yabuki et al. (2013), Telford et al. 2015; Torruella et al. (2015), Whelan et al. (2015), Corsaro et al. (2016), Carr et al. (2017), Dohrmann & Wöhrheide (2017), Hehenberger et al. (2017), Schiffer et al. (2017), Simion et al. (2017), Tedersoo et al. (2018), (Opisthokonta); Yoon et al. (2008, 2011), Wegener Parfrey et al. (2010, 2011), Burki et al. (2012, 2016), Cavalier-Smith & Chao (2012), Yabuki et al. (2012, 2014) Cavalier-Smith et al. (2014, 2015a), Burki (2014), Katz & Grant (2014), Sharpe et al. (2015), Brown et al. (2017) (minor groups and all eukaryotes). Following the divergence time estimates of Wegener Parfrey et al. (2011), kingdoms and phyla were assigned to higher taxa that diverged roughly at >1000 and 542 Mya (as described for fungi in Tedersoo et al. 2018). These criteria were used to make the latest diverging kingdoms Metazoa and Viridiplantae comparable to other eukaryote groups. Kingdoms that formed well-supported monophyletic groups were further assigned to subdomains. For kingdoms and phyla, I proposed to use currently accepted names, prioritizing widely used names and those referring to particular taxa, which is in line with the zoological and botanical nomenclature. The few newly proposed names are derived from the names of type taxa. Comparisons between classifications are mostly performed against that of Ruggiero et al. (2015), which is the most widely followed and cited (in both positive and negative sense) recent treatment.

## Results and Discussion

### General patterns

Out of multiple classifications, the NCBI and SILVA classifications were the most updated in terms of state-of-the-art phylogenetic information. Compared with the SILVA classification, the NCBI system comprised much less putative names and codes of undescribed taxa, or these were more comprehensively classified into the Linnaean taxonomic framework. Therefore, the NCBI system (as of 12 October 2017) was selected as a baseline for further work.

Based on multiple molecular phylogenies, the monophyly criterion and roughly comparable divergence time, 32 kingdom-level groups were recovered (Figure 2). Most of these were treated at the level of class (in Ruggiero et al. 2015; see Figure 3) or at the level of phylum or no rank (in NCBI). Monophyletic kingdoms were further grouped into 10 subdomains or subdomain-level taxa, of which four (Archaeplastida, Excavata, Harosa and Opisthokonta) are comprised of >1 kingdom (Figure 3). The 32 kingdoms were further divided into 43 subkingdoms (including 14 named), 115 phyla (102), 140 subphyla (51) and 353 classes (305). The classification down to class level and genus level is given in Appendix 1 and supplementary document (http://dx.doi.org/10.15156/BIO/587483), respectively. The lower proportion of named taxa at the level of subranks indicates that subranks were not effectively used in most groups and were left as ‘unspecified’ if monotypic. In relatively well-studied and morphologically diverse kingdoms such as Metazoa, Viridiplantae and Fungi, subkingdoms and subphyla were commonly used to provide more natural grouping and improve resolution.

**Figure 2.**
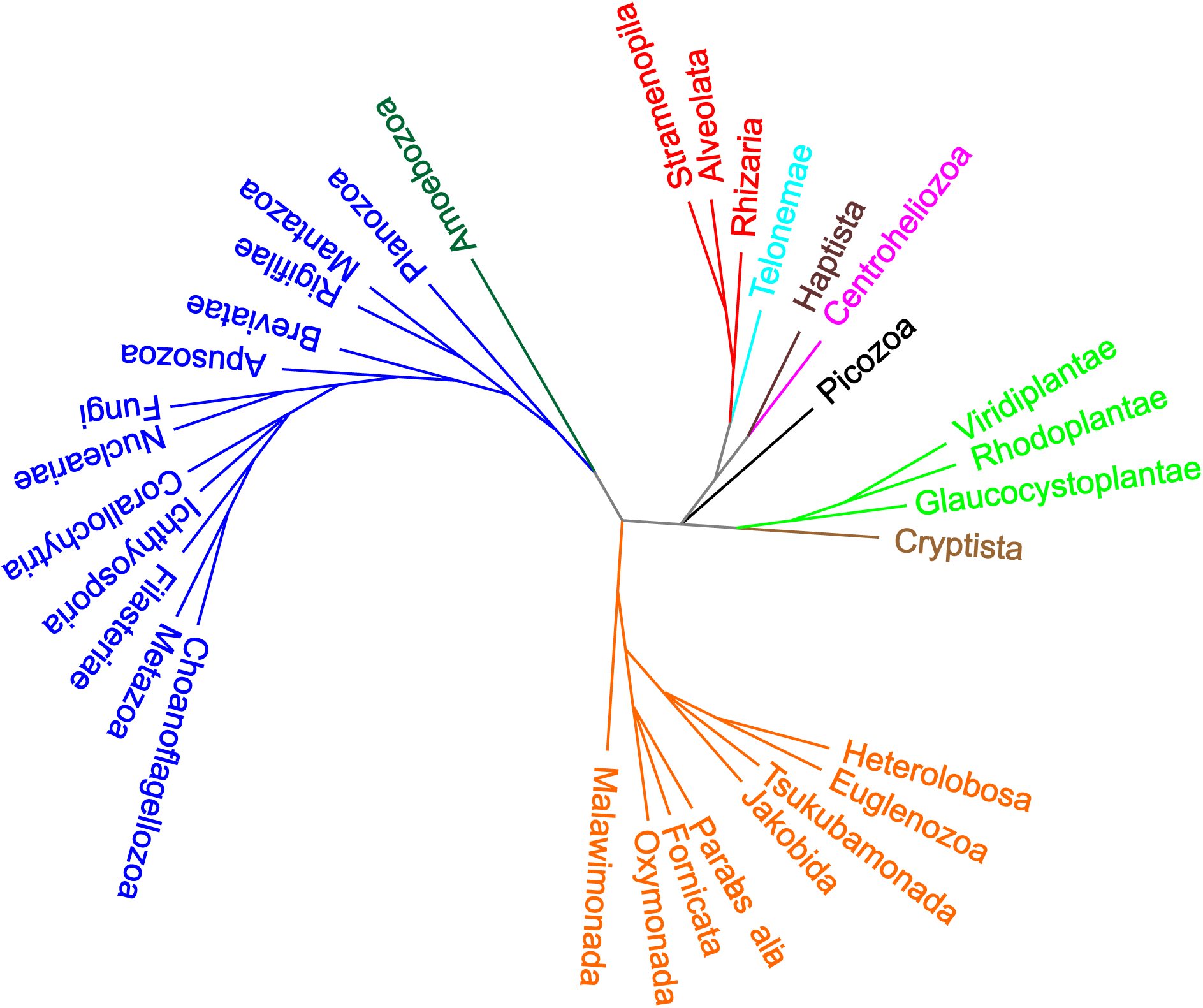
Unrooted taxon tree indicating the proposed kingdom-level classification of the Eukarya domain. Different colours indicate subdomains.

**Figure 3.**
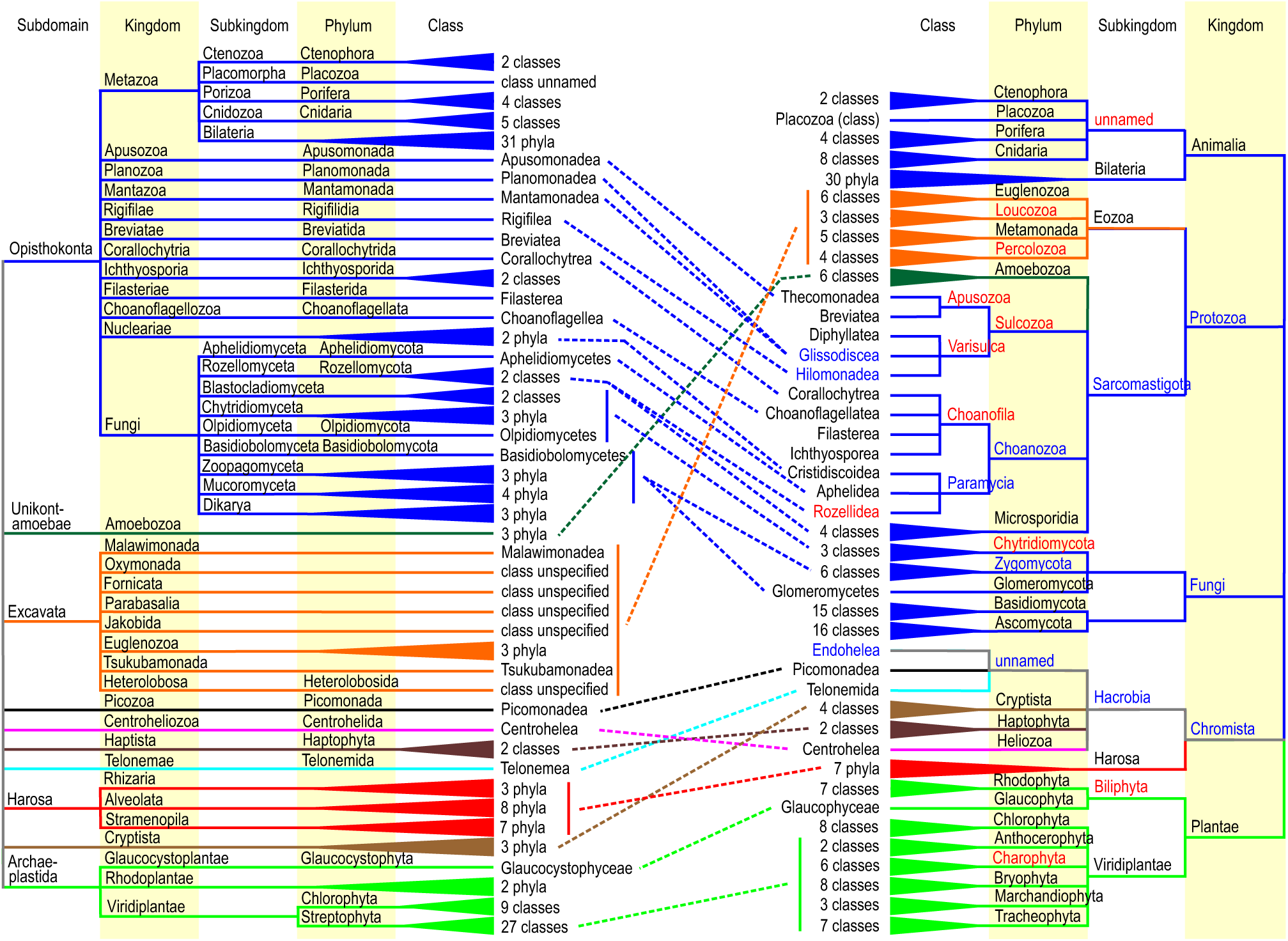
Comparison of higher level classification of Ruggiero et al. (2015; right pane) and the proposed classification (left pane). Red and blue fonts indicate paraphyletic and polyphyletic taxa, respectively; different branch colours depict subdomains; dashed lines indicate corresponding taxon names that differ between classifications.

Compared with Ruggiero et al. (2015) and Adl et al. (2012), the proposed classification contains no taxa that are intentionally erected as paraphyletic. Nonetheless, it is likely that some of the taxa will turn out to be paraphyletic in more refined phylogenomic analyses. In Ruggiero et al. (2015), several taxa in each taxonomic level seem to have been treated as trash bins to accumulate orphan taxa. For example, the phylum ‘Choanozoa’ within kingdom ‘Protozoa’ includes multiple Opisthokontan protists of very different origin including ‘ Aphelidea’ that belong to Fungi. Similarly, an ‘unnamed’ hacrobian phylum within the kingdom ‘Chromista’ includes several classes that are so deeply diverging that these warrant a subdomain and kingdom of their own. Furthermore, ‘Zygomycota’ within Fungi comprises multiple phyla of early diverging mycelial lineages.

### Major subdomains and kingdoms

The subdomain Opisthokonta has been interpreted differently in recent phylogenetic and classification studies by comprising only groups intimately related to Metazoa and Fungi or additionally including all minor deeply diverging taxa that diverged after the Amoebozoa (i.e., ‘Obazoa’). For practical reasons, I recommend to use the broader interpretation for Opisthokonta, because the branching order of smaller groups in not fully settled, to minimize the number of subdomain-level taxa, and use a widely known name. Because the formerly proposed ‘Apusozoa’, ‘Choanozoa’ and ‘Sulcozoa’ are strongly para- or polyphyletic, several minor deeply diverging groups were separated from these to represent distinct kingdoms (i.e., Apusozoa *s.stricto,* Breviatae, Choanoflagellozoa, Corallochytria, Filasteriae, Ichthyosporida, Mantazoa, Planozoa, Rigifiliae). Certain earlier studies indicated that Opisthokonta in the wide sense may be paraphyletic with respect to Amoebozoa, but this is not supported in more recent and more inclusive studies (but see Brown et al. 2017). In Metazoa, it would be feasible to provide a subkingdom-level separation to Bilateria, Ctenophora (as Ctenozoa), Porifera (Porizoa) and Placozoa (Placomorpha). On the fungal side of Opisthokonta, I propose to consider Fungi and Nucleariae (‘Cristidiscoidea’) as distinct kingdoms and recommend acceptance of nine subkingdoms within Fungi. These include Rozellomyceta (including Microsporidea) and Aphelidiomyceta that are closely related to other fungal groups (James et al. 2013; Corsaro et al. 2014; Tedersoo et al. 2018) rather than forming a cluster of their own (‘Opisthosporidia’ hypothesis; Karpov et al. 2014, 2017; Torruella et al. 2017).

The subdomain Unikontamoebae *nom. provis.* (named as such to secure no overlap with kingdom name) is comprised of the kingdom Amoebozoa and forms a coherent, well-supported sister group to Opisthokonta. Amoebozoa is comprised of three phyla (Discosida, Evosida and Tubulinida) following a recent phylogenomic analysis (Tekle & Wood 2017), which challenges the traditional split of Amoebozoa into Lobosa and Conosa. Based on the NCBI classification and published phylogenetic analyses, there is much uncertainty at the level of classes and orders (Cavalier-Smith et al. 2016; Tekle et al. 2016). The seemingly natural group Mycetozoa may be paraphyletic with respect to Archamoebidea and it is separated into two classes - Eumycetozoa and Variosea - within Evosida (Tekle & Wood 2017).

Harosa comprises kingdoms Stramenopila, Alveolata and Rhizaria, collectively known as ‘SAR’ in informal classification (Burki et al. 2007). Both Harosa and the kingdoms therein are phylogenetically well supported. Given their deep divergence, it is recommended to consider the main taxonomic groups in these kingdoms at the phylum level rather than class level. In this treatment, Stramenopila, Alveolata and Rhizaria are comprised of seven, eight and three phyla, respectively, most of which exhibit distinct ecophysiology (Cavalier-Smith 2018). The relatively recently diverged groups Foraminifera, Polycystinea and Acantharea are considered at the class level within the phylum Retaria of Rhizaria. There are several genera that may warrant recognition at the class or phylum level in Alveolata (*Palustrimonas, Voromonas)* and Stramenopila *(Cantina, Pirsonia, Leukarachnion, Platysulcus, Pseudophyllomitus* and multiple ‘MAST’ clades).

The subdomain Excavata has received much less phylogenetic hypothesis testing compared with other major groups. Based on multiple phylogenetic reconstructions, I propose eight kingdoms within Excavata, viz. Heterolobosa, Fornicata, Jakobida, Tsukubamonada, Oxymonada, Parabasalia, Euglenozoa and Malawimonada. Members of the Malawimonada are highly divergent and commonly cluster within or in a sister position to other subkingdoms, but Kolisko (2011) indicated its firm placement within Excavata when removing rapidly evolving regions causing long branches. Although Cavalier-Smith et al. (2014) consider Euglenozoa outside Excavata, other authors have demonstrated its nested position within excavates. Apart from Euglenozoa and Heterolobosa, separation of other excavate kingdoms to phyla was not attempted due to paucity of molecular phylogenetic research. Given the long branches and rapid evolution, the ‘metamonad’ kingdoms Parabasalia, Oxymonada and Fornicata may turn out to be paraphyletic when more detailed information accumulates.

Archaeplastida contains kingdoms Glaucocystoplantae, Rhodoplantae (phyla Rhodophyta and Cyanidiophyta) and Viridiplantae, which is separated into Chlorophyta and Streptophyta at the phylum level. Streptophyta is further divided into multiple subphyla, including groups representing green algae, early land plants and vascular plants – Tracheophytina. The latter is comprised of major fern and gymnosperm groups and Angiospermae at the class level. This class-level treatment represents re-organisation of major plant taxa by the level of rank or subrank due to relatively recent divergence of these plant groups relative to other kingdoms (e.g. Wegener Parfrey et al. 2011).

### Minor kingdoms and unplaced taxa

Many studies place Haptista and particularly Cryptista in a sister position to Archaeplastida, but in most studies these groups branch off separately. Haptista contains two classes (Coccolithophyceae and Pavlovophyceae), which may warrant phylum-level separation. Cryptista is comprised of three deeply diverging phyla (Figure 3). Telonemae is a small groups comprised of two sequenced species (*Telonema antarctica* and *T. subtile*) but represented by a single species in most analyses. Telonemae is most commonly placed in a sister position to the subdomain Harosa (e.g. Cavalier-Smith & Chao 2010; Burki et al. 2012). Picozoa (‘picobiliphytes’) is represented by a single described species *(Picomonas judraskeda)*, although the cryptic diversity of closely related marine taxa is much higher. Picozoa usually branches off separately from any major kingdom or occur in a sister position to Cryptista or Centroheliozoa (Yoon et al. 2011; Moreira & Lopez-Garcia 2014). Centroheliozoa is also known as ‘Heliozoa’, but the latter name has multiple meanings. This kingdom is comprised of 11 sequenced genera that form a coherent group with a long stem (Cavalier-Smith & von der Heyden 2007; Cavalier-Smith & Chao 2012), which warrants treatment of all taxa in a single class Centrohelea, in agreement with Ruggiero et al. (2015). This group has remarkable cryptic diversity in both saltwater and freshwater and soil habitats (Cavalier-Smith & Chao 2012).

There are a few deeply diverging taxa that cannot be reliably related to any proposed subdomain and kingdom (Appendix 1; http://dx.doi.org/10.15156/BIO/587483). This is at least partly ascribed to their inclusion in only a few analyses and/or based on 1–2 genes. *Collodictyon triciliatum* and *Diphylleia rotans* (phylum Collodictyonida) are placed within the class ‘Endohelea’ of hacrobian ‘Chromista’ together with ‘Heliomonadida’ (sensu Ruggiero et al. 2015). The latter order is located in ‘Granofilosea’ of harosean ‘Chromista’ in Bass et al. (2009) and AlgaeBASE (www.algaebase.org). While reliable sequences of Heliomonadida are still unavailable, Collodictyonida is phylogenetically placed in a sister position to Opisthokonta, Excavata or Amoebozoa or other minor groups with long branches (Zhao et al. 2012; Cavalier-Smith et al. 2014). The most recent and analyses place it in a sister position to the phylum Rigifilidia within Opisthokonta (Brown et al. 2017). *Microheliella maris* (phylum Microheliellida) is phylogenetically distinct from other taxa, but may have some affinities to Centroheliozoa, Telonemae or Cryptista (Cavalier-Smith & Chao 2012; Yabuki et al. 2012).

### Conclusions and perspectives

Single-cell genomics and trancriptomics methods and phylogenomics analyses have become available in the last 8 years and enabled to resolve the order to phylum level internal structure in many kingdoms. However, these methods still lack sufficient power to provide reliable placement of the minor kingdoms and highly divergent obligately parasitic or anaerobic taxa that are represented by 1–2 isolates. At this stage, certainly more diversity in these groups must be captured, which is of great importance to be able to understand the entire eukaryote evolution (Pawlowski 2013). In unculturable organisms, single cells for genomics analyses can be obtained by using fluorescent probes specifically targeting their DNA (e.g. Jones et al. 2011). It can be speculated that multiple novel kingdoms and phyla are yet to be recovered, which has been demonstrated for many groups. However, as pointed out by Berney et al. (2004), many of these supposedly novel groups represent chimeric marker sequences or rapidly evolving taxa that find their position in more inclusive analyses. Out of >40 novel soil fungal groups, three have been described or matched to sequenced specimens 24 months after analysis (Tedersoo et al. 2017).

The proposed classification of eukaryotes represents a consensus of multiple phylogenetic studies, which is based on the monophyly criterion and rough divergence time estimates (Avise & John 1999). The other four typical problems in modern classification systems (Figure 1) were accounted for as much as possible, but their handling required almost always compromises between selecting appropriate, non-overlapping, well-known names and erection of optimal number of ranks, which was partly influenced by previous use of these names and availability of phylogenetic information. The proposed names are usually directly linkable to the type genus or forms of these have been widely accepted by the research community. My team also proposed a ‘taxon hypothesis’ concept to be able to cross-link different classification systems in space and time (Tedersoo et al. 2018).

The proposal of multiple kingdoms is not new in science. For example, in pre-molecular era, Leedale (1977) presented a classification system with 18 eukaryote kingdoms, most of which represented various protist groups and early diverging Metazoa. Compared with the classification presented here, Leedale’s kingdoms included taxa from the level of class to subdomain. His treatment certainly represented state-of-the-art of the contemporary knowledge, but the system was not accepted by the scientific community. Most recently, Drozdov (2017) proposed a classification including 15 eukaryote kingdoms. However, this treatment is hardly comparable to other modern classifications, because it is a mixture of morphological and phylogenetic classifications that forces several class-level groups such as Foraminifera and Microsporidea at the kingdom level, and keeps most kingdoms paraphyletic.

In the proposed classification, the erection of 32 eukaryote kingdoms certainly catches and, perhaps, scratches the eye. I found adoption of multiple kingdoms necessary to follow the monophyly principle and to render the relatively late diverging kingdoms Metazoa and Viridiplantae better comparable to multiple protist groups in terms of divergence time. To be strict, it might still require lumping the Choanoflagellozoa to its sister group Metazoa and splitting Unikontamoebae and Excavata to additional kingdoms. Such activities may require revising the age estimates of major eukaryote groups based on additional calibration points and genomic comparisons.

Looking ahead, such multiple-kingdom classification approach would tremendously improve taxonomic resolution of Bacteria and Archaea, for which the kingdom rank is essentially unused (Woese et al. 1990). In contrast to other classifications, Drozdov (2017) erected four and seven kingdoms to accommodate phyla of Archaea and Bacteria, respectively, but most of the proposed groups are para- or polyphyletic. Since prokaryotes evolved and diverged >3 billion years ago (Sheridan et al. 2003), their classification might require another rank between kingdom and domain (for example, rejuvenating the *empire* rank) to accommodate the earliest branching clades. If monophyletic, all prokaryote groups hitherto recognized at the phylum level could be instantly ascribed to separate kingdoms given their time of divergence and improved comparability to eukaryotes

Taken together, I advocate that modernizing the classification of life is necessary for ease of communication between taxonomists, ecologists and molecular biologists. The criteria of monophyly, roughly comparable divergence times and names deduced from genus names are likely to render the names of higher-level taxa much more long-lived and acceptable to the scientific community. I hope that this preprint raises a heavy discussion among taxonomists and leads the way to a modern, global classification system of all life.

## Acknowledgements

I thank K. Põldmaa for constructive comments on a pre-submission version of the manuscript. I received funds from Estonian Science Foundation grant PUT1399, MOBERC10 and Centre of Excellence ECOLCHANGE.

---

**Appendix 1.**
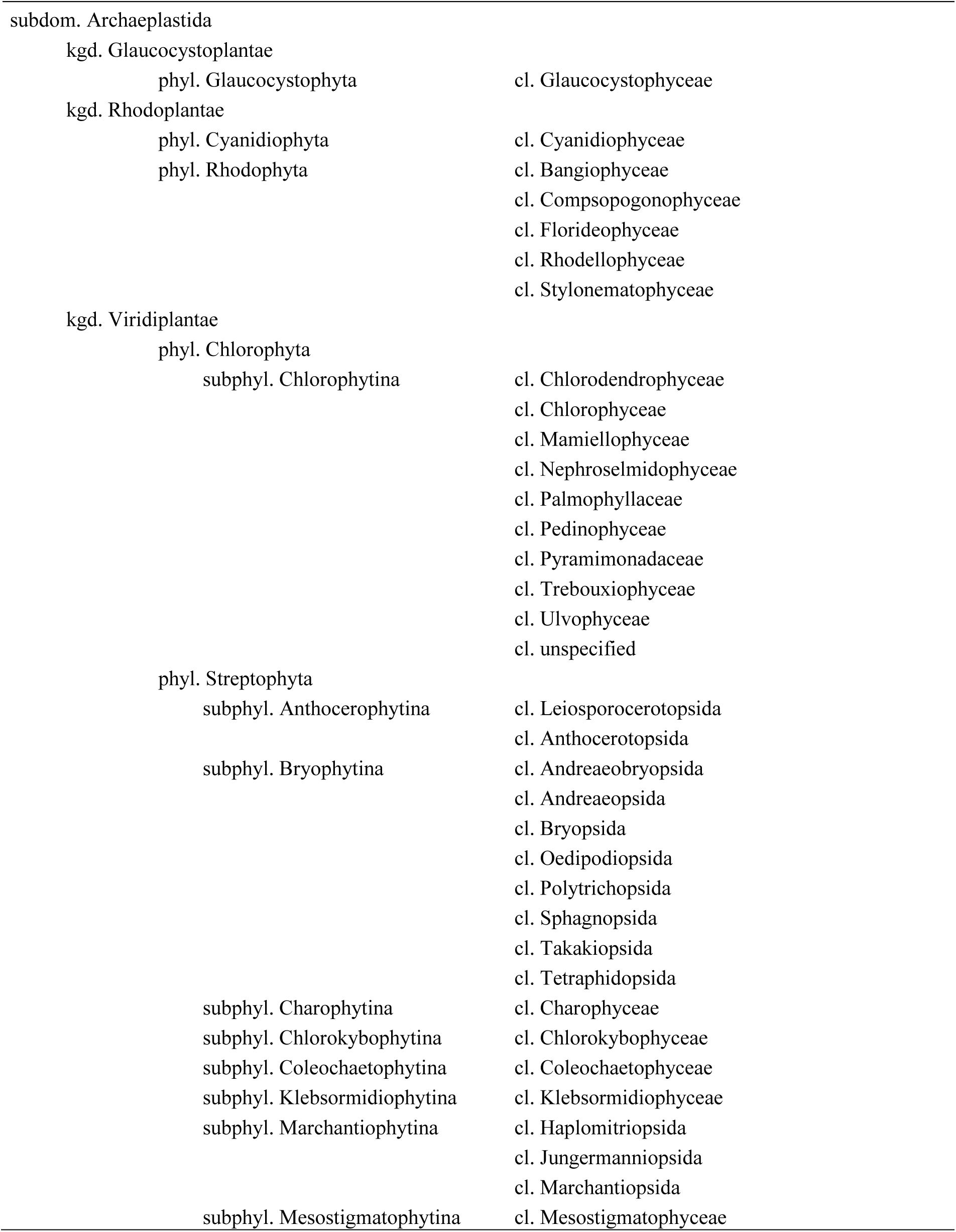
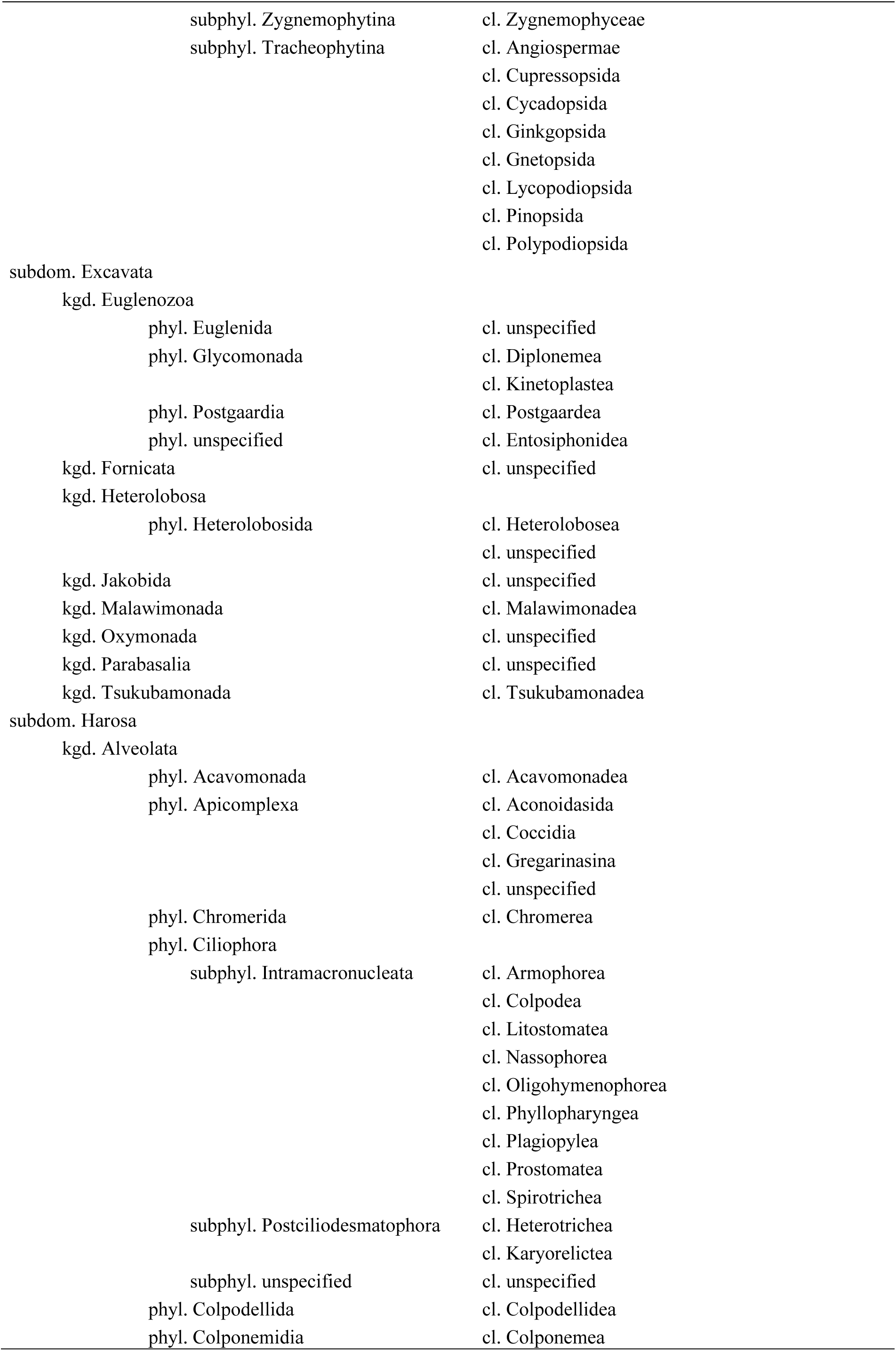
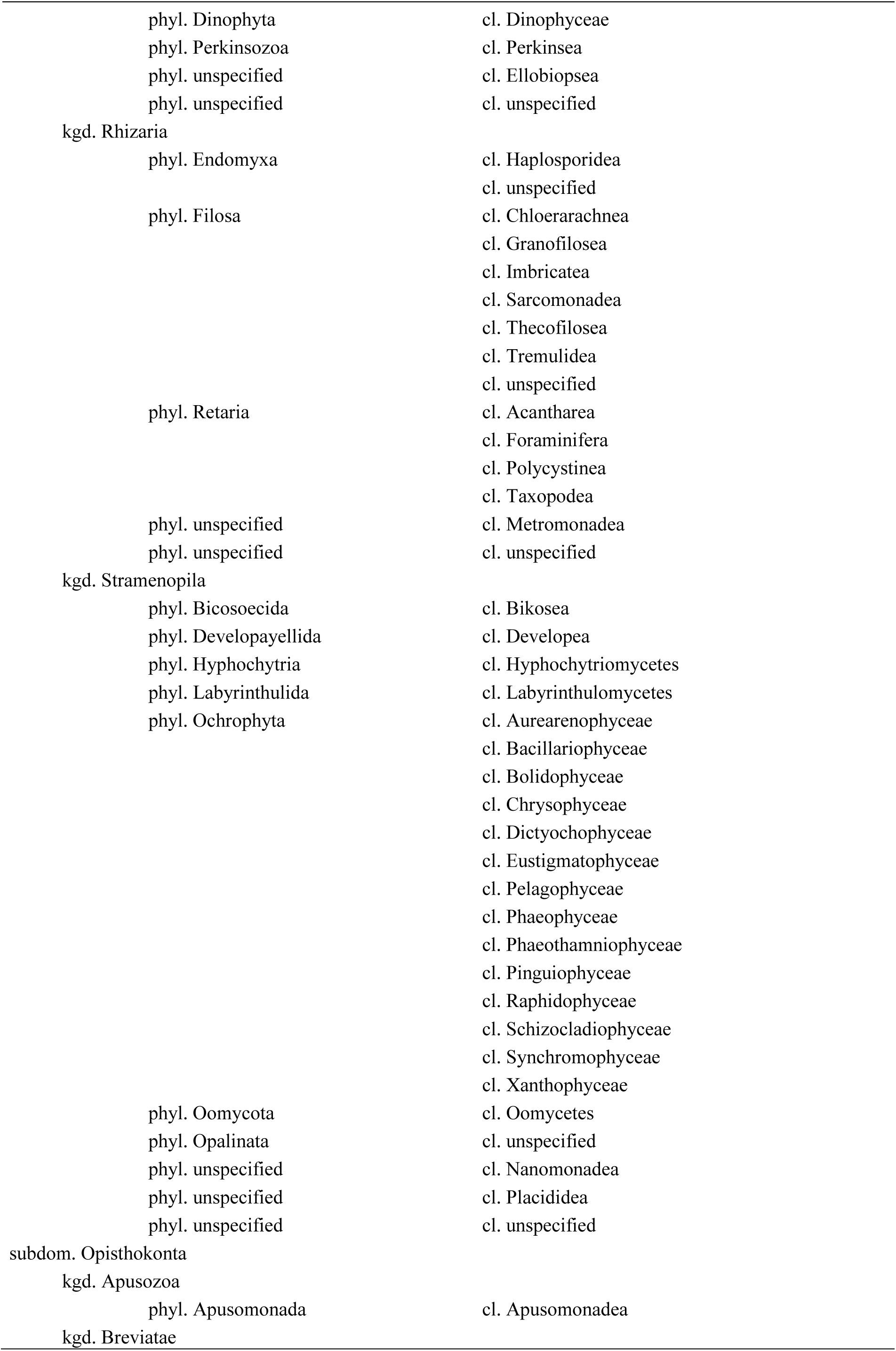
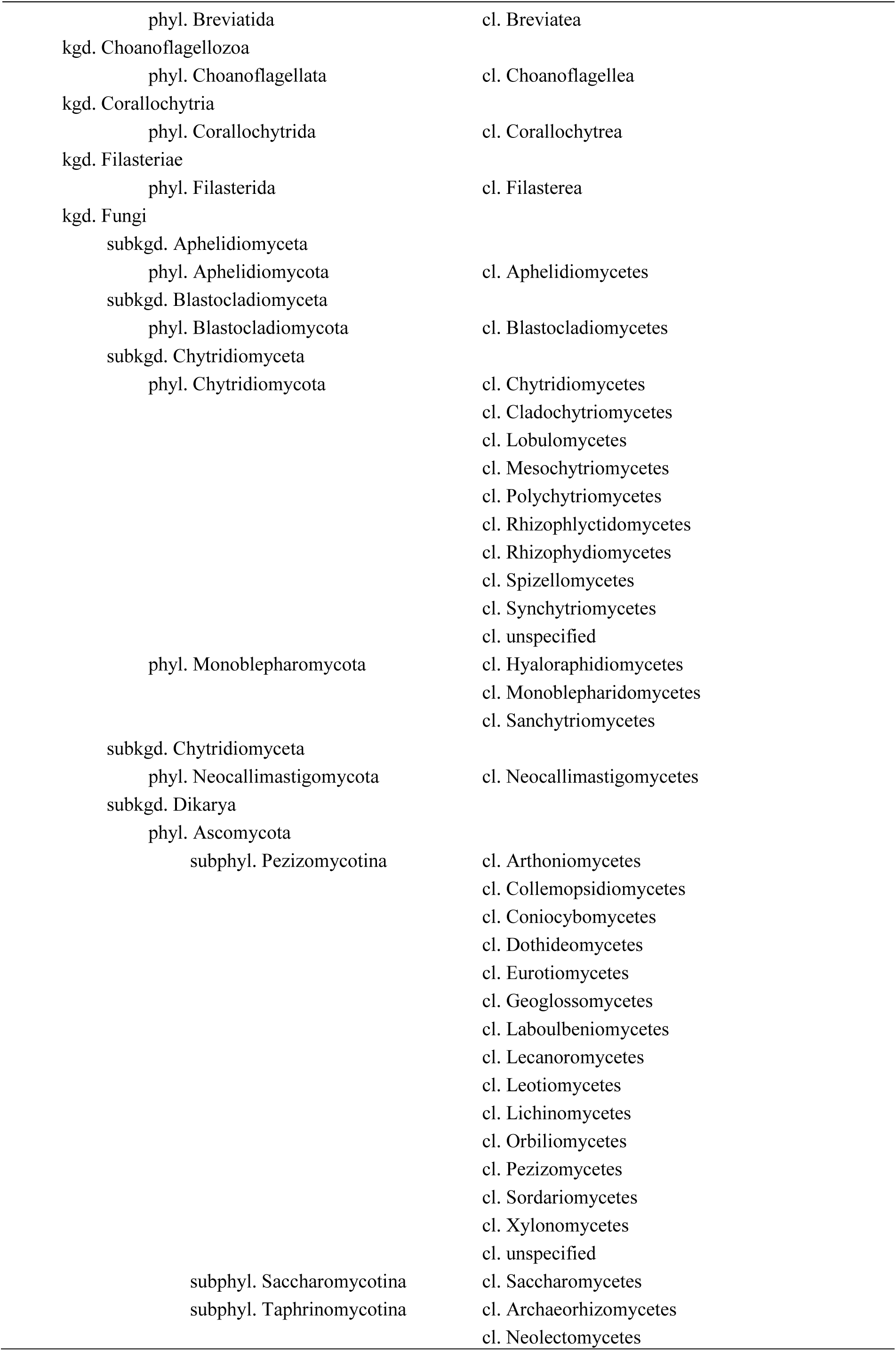
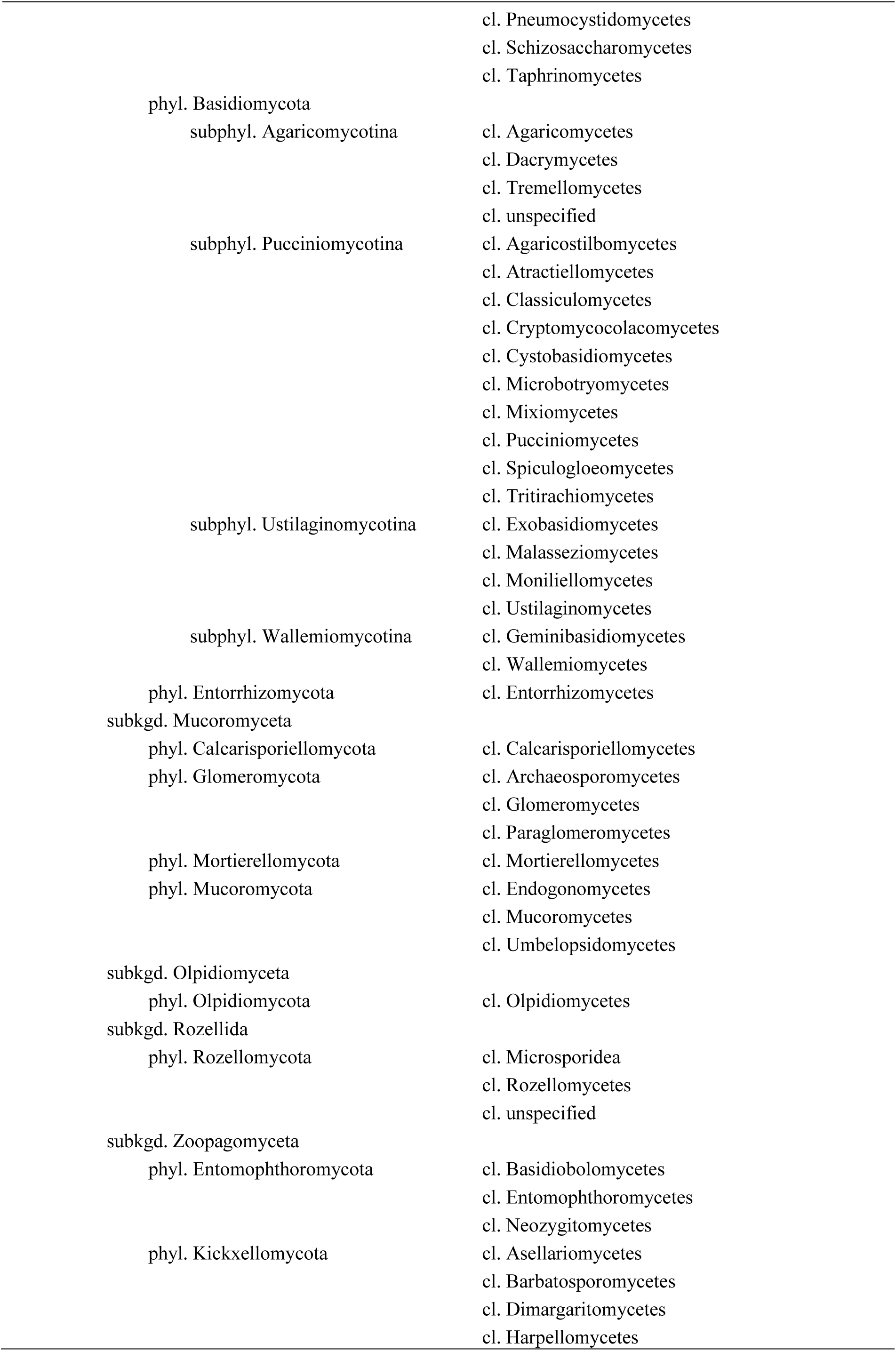
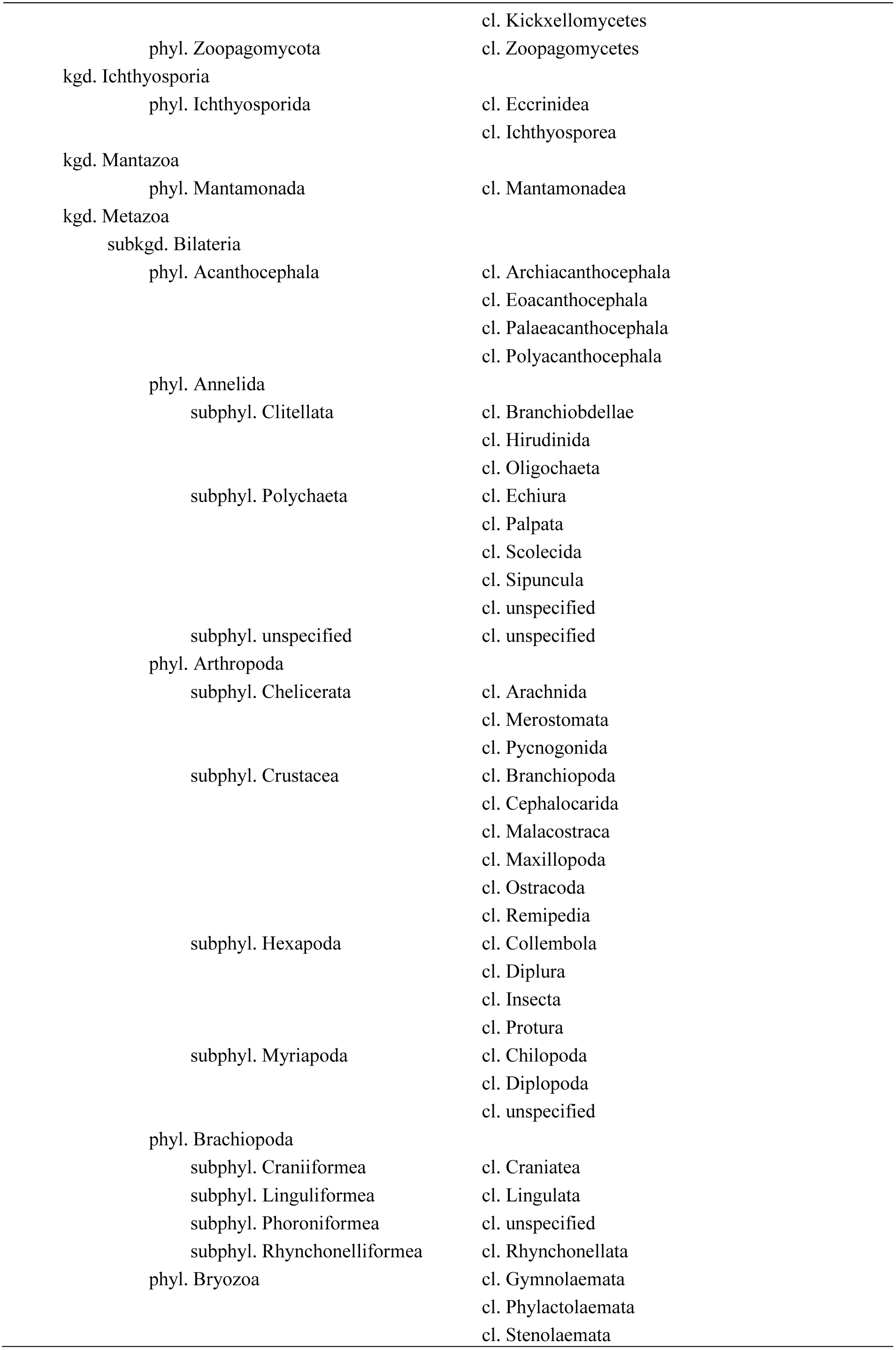
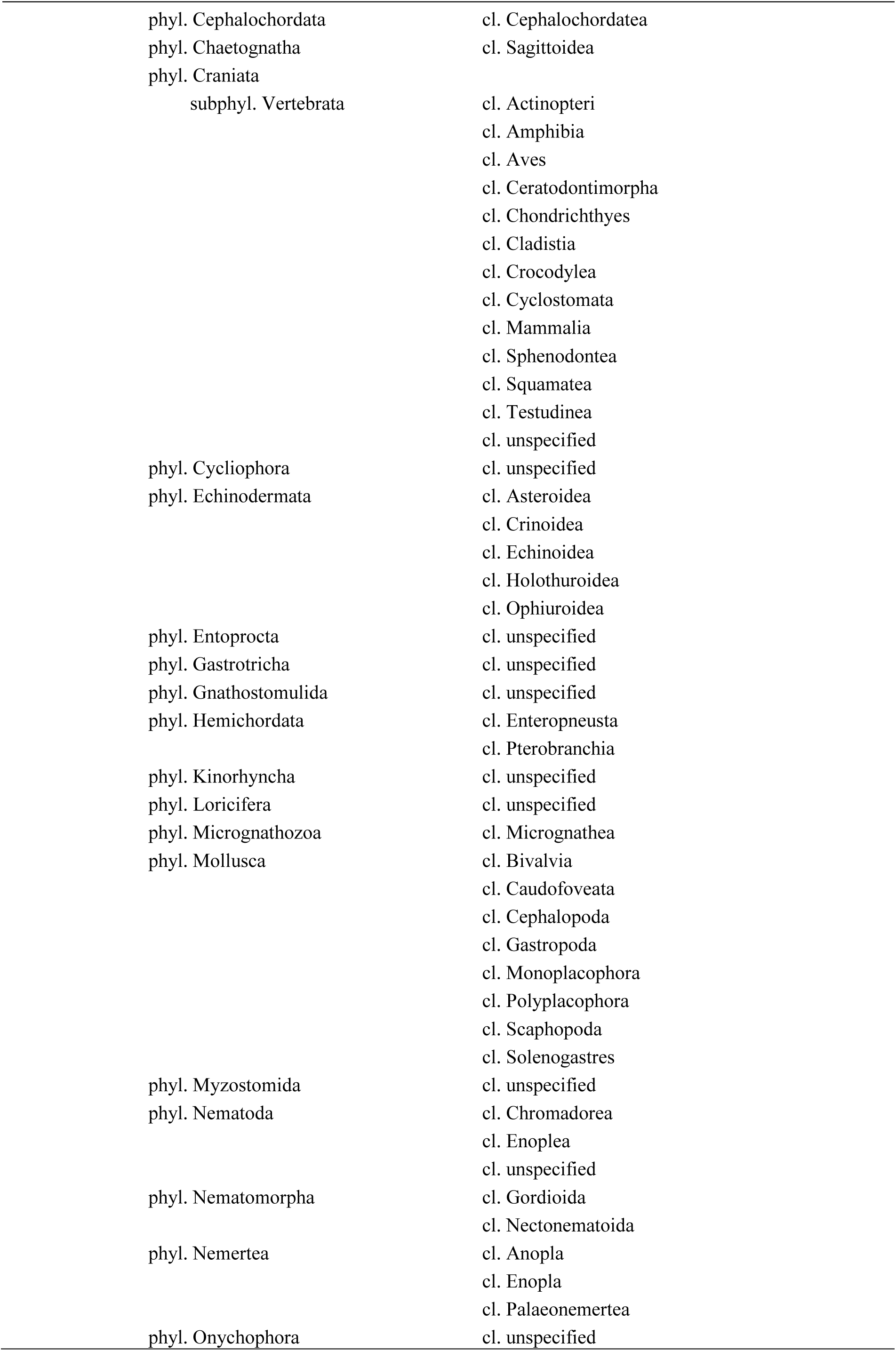
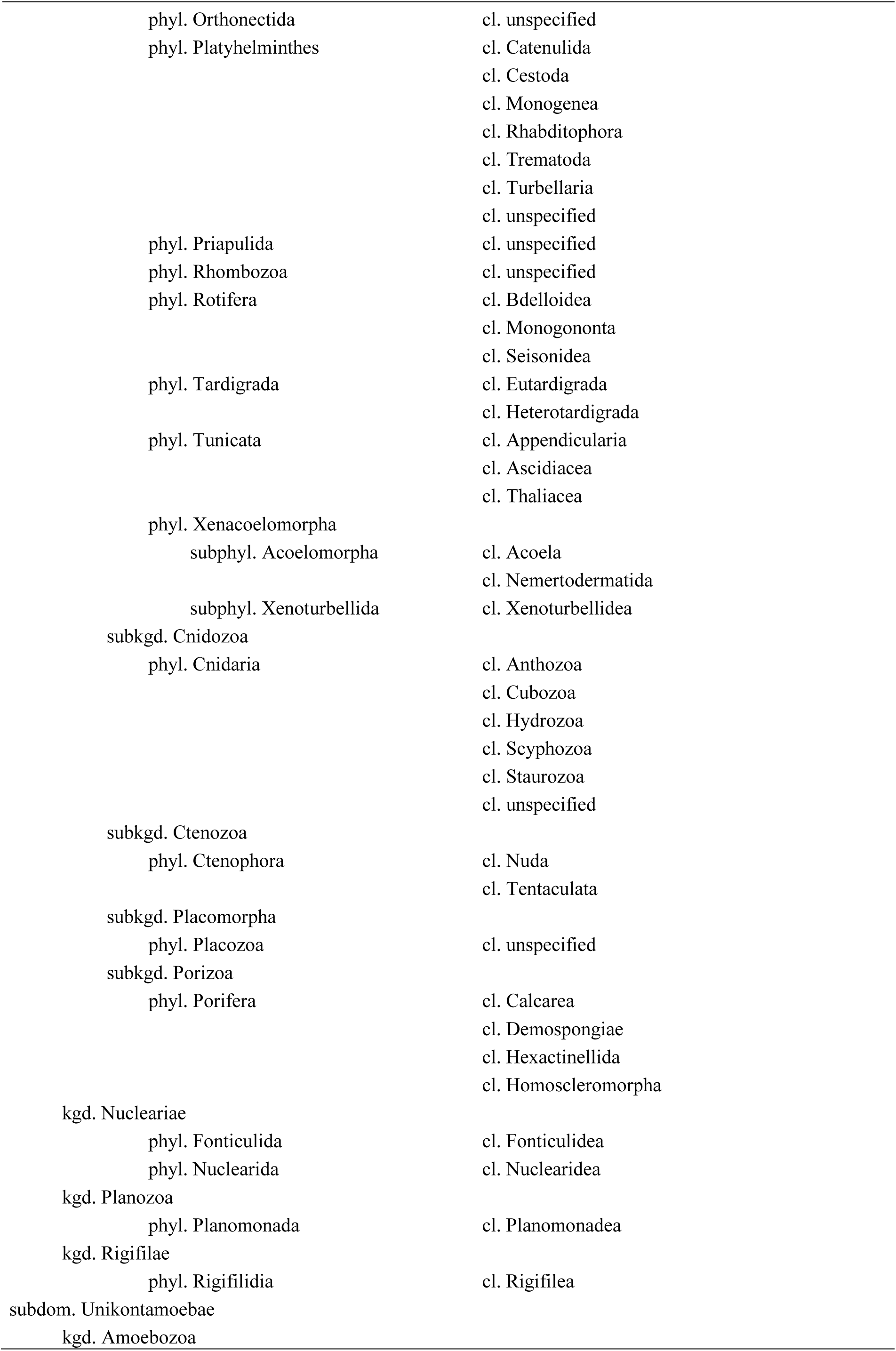
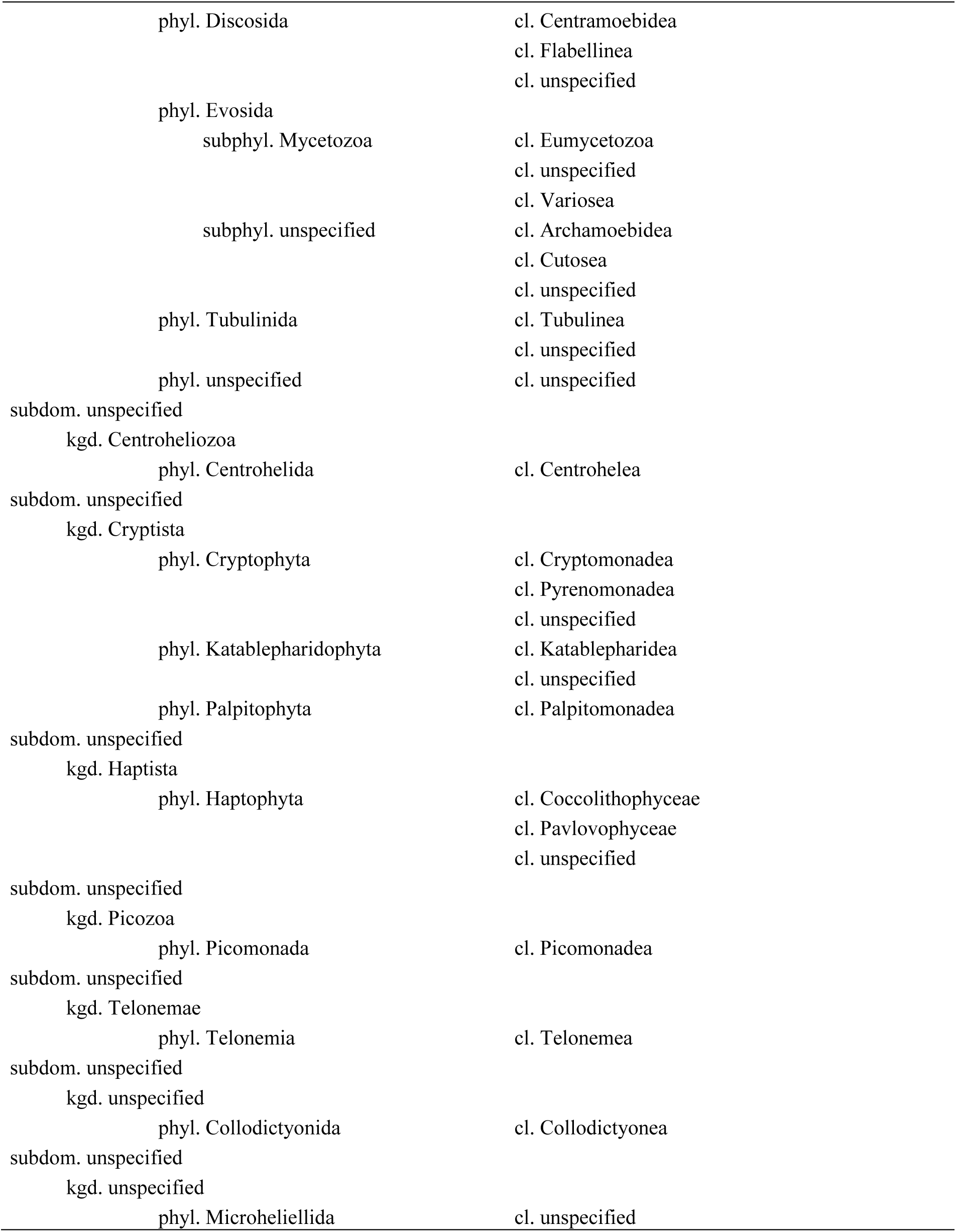
Subdomain to subphylum level classification of eukaryotes with ingredient classes indicated. Other groups not described at particular taxonomic level are included as ‘unspecified’ taxa. The full NCBI-based classification table down to genus level is given in the associated material (http://dx.doi.org/10.15156/BIO/587483)

**Associated material.** Tedersoo L. 2017. Proposed practical classification of the domain Eukarya based on the NCBI system and monophyly and comparable divergence time criteria. (http://dx.doi.org/10.15156/BIO/587483)

